# Zika virus-like particles bearing covalent dimer of envelope protein protect mice from lethal challenge

**DOI:** 10.1101/2020.07.09.196626

**Authors:** Giuditta De Lorenzo, Rapeepat Tandavanitj, Jennifer Doig, Chayanee Setthapramote, Monica Poggianella, Ricardo Sanchez Velazquez, Hannah E. Scales, Julia M. Edgar, Alain Kohl, James Brewer, Oscar R. Burrone, Arvind H. Patel

## Abstract

Zika virus (ZIKV) envelope (E) protein is the major target of neutralizing antibodies in infected host, and thus represents a candidate of interest for vaccine design. However, a major concern in the development of vaccines against ZIKV and the related dengue virus is the induction of cross-reactive poorly neutralizing antibodies that can cause antibody-dependent enhancement (ADE) of infection. This risk necessitates particular care in vaccine design. Specifically, the engineered immunogens should have their cross-reactive epitopes masked, and they should be optimized for eliciting virus-specific strongly neutralizing antibodies upon vaccination. Here, we developed ZIKV subunit- and virus-like particle (VLP)-based vaccines displaying E in its wild type form, or E locked in a covalently linked dimeric (cvD) conformation to enhance the exposure of E dimers to the immune system. Compared with their wild-type derivatives, cvD immunogens elicited antibody with higher capacity of neutralizing virus infection of cultured cells. More importantly, these immunogens protected animals from lethal challenge with both the African and Asian lineages of ZIKV, impairing virus dissemination to brain and sexual organs. Moreover, the locked conformation of E reduced the exposure of epitopes recognized by cross-reactive antibodies and therefore showed a lower potential to induce ADE *in vitro*. Our data demonstrated a higher efficacy of the VLPs in comparison with the soluble dimer and support VLP-cvD as a promising ZIKV vaccine.

**Author Summary:** Infection with Zika virus (ZIKV) leads to the production by host of antibodies that target the viral surface envelope (E) protein. A subset of these antibodies can inhibit virus infection, thus making E as a suitable candidate for the development of vaccine against the virus. However, the anti-ZIKV E antibodies can cross-react with the E protein of the related dengue virus on account of the high level of similarity exhibited by the two viral proteins. Such a scenario may lead to severe dengue disease. Therefore, the design of a ZIKV vaccine requires particular care. Here, we tested two candidate vaccines containing a recombinant form of the ZIKV E protein that is forced in a covalently stable dimeric conformation (cvD). They were generated with an explicit aim to reduce the exposure of the cross-reactive epitopes. One vaccine is composed of a soluble form of the E protein (sE-cvD), the other is a more complex virus-like particle (VLP-cvD). We used the two candidate vaccines to immunize mice and later infected with ZIKV. The animals produced high level of inhibitory antibodies and were protected from the infection. The VLP-cvD was the most effective and we believe it represents a promising ZIKV vaccine candidate.

## Introduction

For decades Zika virus (ZIKV) was largely ignored as a human pathogen but the recent epidemic in South America has brought to light neurological complications (i.e. Guillain-Barré syndrome)(1) and congenital Zika syndrome (i.e. microcephaly and other malformations)(2) making ZIKV a public health threat in affected countries. ZIKV infection occurs mainly via mosquito bite but its persistence in bodily fluids like semen allows sexual transmission (3). There is currently no vaccine or treatment available, making their development a priority in ZIKV research.

Current approaches to vaccine development include purified inactivated virus (4), DNA/RNA/vector-based vaccines encoding structural proteins (5-10) and purified viral-like particles (VLPs) (11, 12) or protein subunits (13, 14). Some of these candidates are currently undergoing phase 1 of clinical trial but the design of a successful ZIKV vaccine is complicated by the close relation of ZIKV with other flaviviruses and especially dengue virus (DENV), also transmitted by *Aedes* mosquito vectors and overlapping across many areas.

ZIKV genome, like other members of the *Flaviviridae* family, is composed of a positive strand RNA encoding a single polyprotein that is cleaved into structural (capsid, precursor-membrane and envelope) and non-structural proteins (NS1, NS2, NS3, NS4 and NS5). The envelope (E) glycoprotein, with its three domains (DI, DII and DIII), is the main target of the host immune response (15). During the initial stages of flavivirus genesis, the E protein is associated with the precursor-membrane protein (prM) and assumes a trimeric conformation; only during the passage in the trans-Golgi network, where the viral particle encounters an acidic environment, the trimers dissociate to re-assemble as dimers (16). This new conformation is necessary to allow furin-mediated cleavage of prM into pr and M generating a mature E dimer (17). Once released in the extracellular environment, pr dissociates and the particle becomes infectious. During infection, the low pH of the endosome triggers a new conformational modification that mediates fusion of viral and endosomal membranes (18). However, the particle maturation process is often incomplete releasing a viral progeny partially displaying E protein in trimers containing prM. In addition, E protein is in continuous dynamic motion, a phenomenon called “virus breathing” that is strain- and temperature-dependent (19). These two factors - incomplete maturation and viral breathing - have important consequences on epitope accessibility. At its tip, DII harbours the fusion loop (FL) represented by an amino acid sequence that is highly conserved among flaviviruses. FL is masked by DI and DIII when E protein on the virion is in a dimeric form but becomes exposed upon re-arrangement of E in the acidic endosome following cell entry. Epitopes located on DI/DII, especially in the FL region (FLE), are immuno-dominant but recognized by cross-reactive and poorly neutralizing antibodies (20, 21). This class of antibodies can be responsible for antibody-dependent enhancement (ADE) of infection where antibody-bound virus particles are endocytosed via the Fcγ receptor, leading to a more severe infection (22). Antibodies to prM also contribute to ADE (22).

In addition, the most potent neutralizing antibodies often recognize complex quaternary epitopes than bind to multiple adjacent E proteins, epitopes that are available only when the E protein is assembled in a viral particle and therefore could not be elicited upon immunization with subunits (23). Recently, a new class of quaternary epitopes, called the Envelope Dimer Epitopes (EDE), have been described (24). EDE epitopes are displayed when the E proteins form a head-to-tail dimeric conformation. Highly neutralizing antibodies recognizing EDE were discovered in the sera of DENV-infected patients but interestingly, they were also shown to efficiently neutralize ZIKV, both in *in vitro* and in *in vivo* experiments (25-27). Upon binding to pre-fusion E dimers, these antibodies can prevent the transition of E to a trimeric form and consequently abrogate membrane fusion and infection.

Here, we aimed to develop antigens for ZIKV vaccination that can drive the immune response preferentially against quaternary/complex epitopes, to increase the neutralizing potential. Our recent study demonstrated that the introduction of a disulphide bridge by A264C substitution can stabilize E in a covalent dimer (cvD) conformation (28). This structure reduces the exposure of the unwanted FLE in favour of EDE. We generated cvD forms of a soluble E (sE-cvD) and a virus-like particle (VLP-cvD). The latter is expected to present E predominantly in form of dimers, conferring a smooth surface to the particles. Vaccination of mice with these antigens afforded full protection from lethal ZIKV challenge. Moreover, in comparison to their WT counterparts, the cvD immunogens elicited antibodies that exhibited lower *in vitro* ADE of DENV, yellow fever virus (YFV) and West Nile virus (WNV). Our data confirmed the potential of cvD mutation in generating an immune response against neutralizing conformational epitopes, and further identified VLP-cvD as the most promising candidate of the two cvD derivatives tested.

## Results

### Design, expression and purification of E covalent dimer-based vaccines

We focused on designing antigens that would elicit antibodies to the complex quaternary epitopes that span two or more ZIKV E molecules. Immunogens based on EDE have a great potential, but a stable dimeric conformation of E is not easy to achieve. For this reason, we used a strategy of generating a covalently stable dimeric form by introducing Ala to Cys mutation in DII (A264C) of E as described previously (28). The stable dimeric E generated is thus expected to enhance exposure of EDE and reduce presentation of the unwanted immune-dominant FL region in DII to the immune system.

We generated a V5 epitope-tagged soluble ZIKV E (sE; i.e. E lacking its stem and membrane anchor domains) in its wild-type form (sE-WT), and in the form of a covalently stabilised dimer (sE-cvD) containing the A264C mutation (Fig 1A) (28). In addition, we also generated wild type (WT) and cvD forms of ZIKV virus-like particles (VLPs) using plasmid constructs encoding the capsid anchor region (Ca) followed by the full-length prM and E (Fig 1C). These proteins were produced by transient transfection of Expi293F cells with the relevant constructs and subsequently purified from the cell medium as described in Methods. The purified sE proteins were analysed in SDS-PAGE gel under reducing and non-reducing conditions (Fig 1B). As expected, the sE-WT was visible exclusively in the monomeric form, with an apparent molecular weight of ∼50 kDa, while sE-cvD under non-reducing conditions had an apparent molecular weight corresponding to a dimer (∼110 kDa) that could be reduced to a monomer upon incubation with DTT. A small amount of sE-cvD was seen in a monomeric form under non-reducing condition. VLPs were expressed in a similar fashion and purified as shown in Fig 1C. SDS-PAGE and western blot analysis confirmed the presence of dimeric E in VLP-cvD (∼110 kDa) when analysed under non-reducing conditions, and this was reduced to a monomer in the presence of DTT (Fig 1D). We also observed two additional minor bands: one in the non-reducing gel where a higher molecular weight protein possibly representing a more complex aggregate of E, and an approximately 90 kDa protein in the reducing gel which is likely an intermediate product resulting from an incomplete thiol reduction. In contrast, monomeric E (∼55 kDa) was found in the VLP-WT preparation under both reducing and non-reducing conditions. The molecular weight of E in the VLP preparations was higher than that of the two sE proteins (Fig 1B) on account of the presence of the stem and anchor sequences. As expected, the viral M (10 kDa) was also detected in both forms of VLPs. Protein M is the product of furin-mediated cleavage of prM (25 kDa) during maturation of virus particles. The presence of M protein in the absence of prM suggested that in VLP-cvD the mutated glycoprotein underwent a complete maturation process during its synthesis, yielding smooth particles bearing the cvD E protein. Instead, the VLP-WT preparation contained residual prM implying that they were not fully matured (Fig 1E). Electron micrographs (Fig 1F) of both types of VLPs showed particles of around 50 nm comparable to the size of infectious ZIKV particles (29).

**Figure 1:**
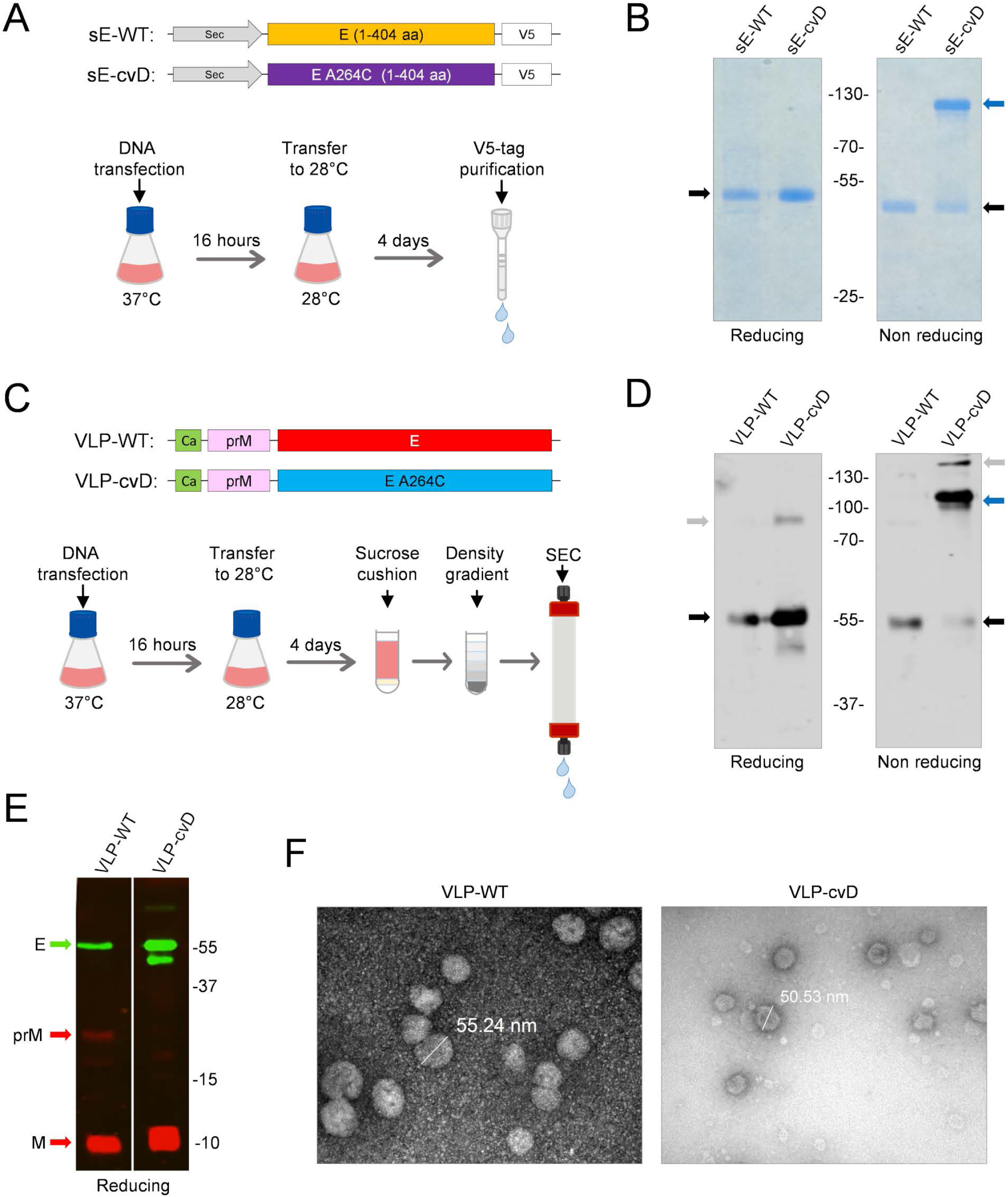
Expression, purification and characterisation of ZIKV E antigens. (**A)** Schematics of the genetic constructs used to express SV5-tagged ZIKV sE (amino acid or aa 1 – 404) in the WT or cvD (carrying A264C mutation) form. Expi293F cells were transiently transfected with the relevant constructs and the expressed proteins secreted into the medium were purified using V5-tag affinity chromatography. (**B) SDS-PAGE of the purified protein.** sE-WT and sE-cvD were analysed in SDS-PAGE run in presence or absence of reducing conditions. Black arrows show monomers of sE, blue arrows show dimers. (**C) Schematics of VLP antigens design and purification:** Plasmid constructs carrying the sequences encoding the capsid anchor (i.e. the N-terminal 18 amino acids of Capsid protein), followed by prM and full-length E genes, the latter in its WT form or in the form of cvD (i.e. carrying the A264C mutation). The constructs were transiently transfected in Expi293F cells and the secreted VLPs were pelleted by sucrose cushion 4 days post-transfection and subsequently purified by density gradient followed by size-exclusion chromatography (SEC). (**D) Western Blot of the purified VLPs showing dimeric conformation:** Purified VLP-WT and VLP-cvD were analysed by SDS-PAGE under reducing or non-reducing conditions E was detected using in-house made monoclonal antibody DIII-1B. Black arrows show E monomers; blue arrow shows dimers; grey arrows show higher order oligomers (non-reducing gel) or partially resolved complexes (reducing gel) of the E protein. **(E) Western Blot of the purified VLPs showing prM and M content:** Purified VLP-WT and VLP-cvD were analysed by SDS-PAGE (14% acrylamide) under reducing conditions. Protein E was detected using the monoclonal antibody DIII-1B (in green), whereas proteins prM and M were detected using an anti-M antibody (in red). (**F) Electron microscope picture of purified VLPs:** Electron microscopy (uranyl acetate staining) of VLP-WT (left panel) and VLP-cvD (right panel) purified as described in (c). Bars indicate the diameter of the particles.

### Antibodies generated by cvD immunogens are conformation-sensitive

ZIKV cannot infect immuno-competent mice due to its inability to counteract murine interferon response (30). We therefore used the interferon receptor-deficient transgenic knock-out (*Ifnar1*^*-/-*^) A129 mice, which are susceptible to ZIKV infection and which have been shown to be amenable to vaccine evaluation studies (31). A cohort each of 4 weeks-old mixed male and female animals (n=6) were vaccinated with sE-WT, sE-cvD, VLP-WT, VLP-cvD, or PBS (as control). Three doses of 10 µg (sEs) or 2 µg (VLPs) of protein adjuvanted with ALUM (1%) combined with MPLA (5 µg) were administered by sub-cutaneous route as shown in Fig 2A. One week after the last dose, blood samples were tested for the presence of anti-E antibodies.

**Figure 2:**
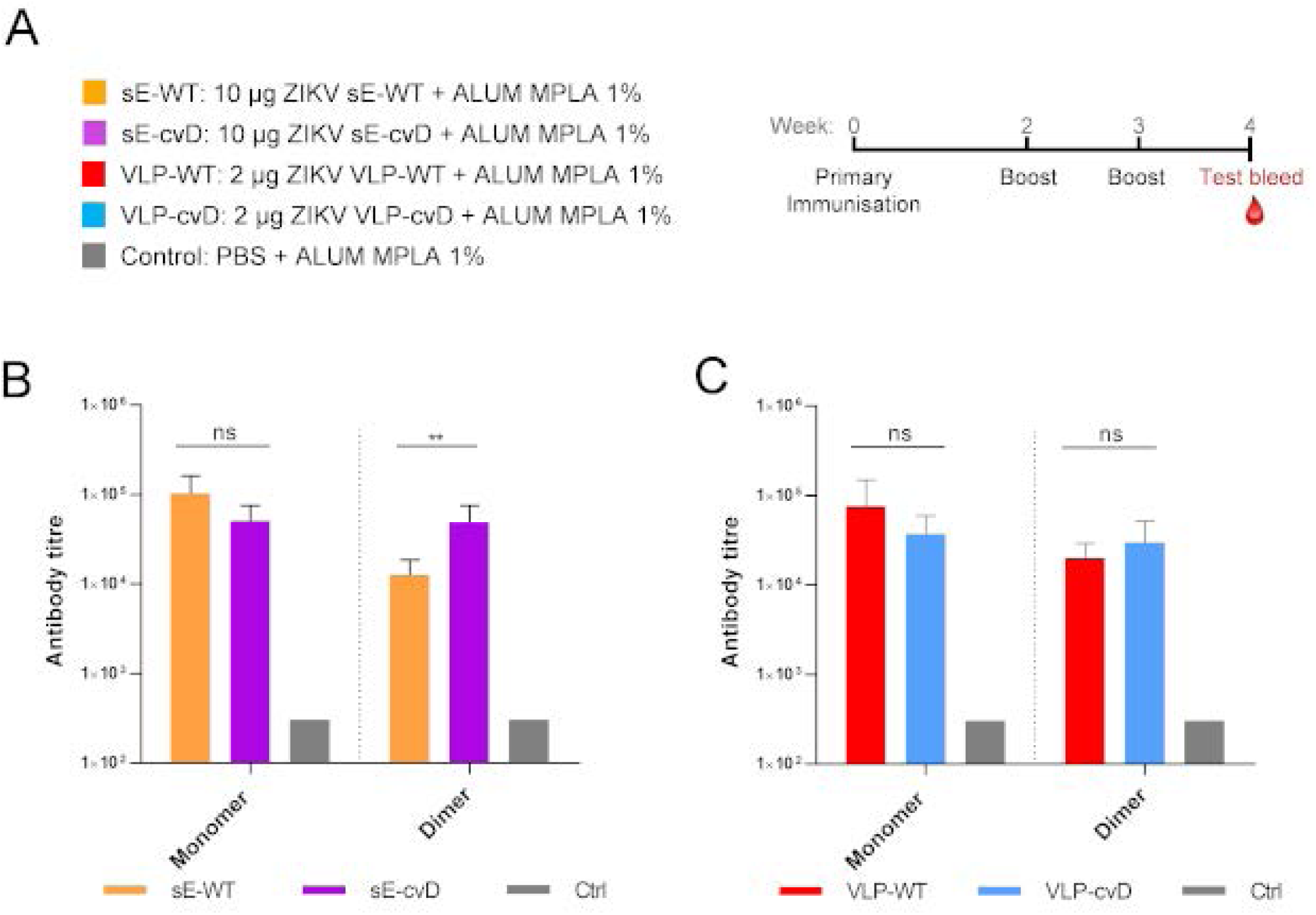
(A) Schematic representation of the immunisation procedure: Five groups each (n=6) of 4 weeks-old mice received respectively sE-WT, sE-cvD, VLP-WT, VLP-cvD or PBS, mixed with ALUM-MPLA adjuvant as shown. Following two boosts at weeks 2 and 3, test bleeds were collected at week 4 for analyses. (**B and C) Anti-E antibody titres of sera collected from animals immunised with sE (B) or VLP proteins (C).** Antibody titres were determined using ELISA plates coated with mono-biotinylated monomeric E, dimeric E and DIII. Ctrl: pooled sera from PBS control group. The titre was defined as the maximum dilution that gives a value higher than three-times the value given by the pre-immune sera. The control sera were negative at the lowest dilution (1:900) and their titre was calculated as 1/3 of that dilution (300). Statistical analysis was done using 2-sided ANOVA 95% confidence level with Tukey Pairwise comparison at 95% confidence (Minitab software).

We first tested the serum antibodies for binding to biotinylated derivatives of ZIKV sE proteins fused in-frame with the Biotin-Acceptor Peptide (BAP). Specifically, recombinant sE-WT-BAP (monomer) or sE-cvD-BAP (dimer) co-expressed with the bacterial biotin ligase BirA (32, 33) (to allow *in vivo* mono-biotinylation) were used to quantify the titre of antibodies recognizing E in its monomeric or dimeric form.

A comparison of the binding levels showed that sE-cvD immunisation elicited antibodies titres against the dimer four times higher than those obtained with sE-WT. On the other hand, analysis of the sera from VLP-WT- and VLP-cvD-vaccinated animals by ELISA showed no significant changes in the levels of antibodies (Fig 2C). It should be noted that this ELISA format is not robust enough to discriminate antibodies binding to more complex epitopes.

In order to further characterise the types of antibodies elicited by our cvD antigens, we used a recently developed cytofluorimetry assay of cells displaying dimers of ZIKV sE protein (paper in submission). In this assay, the C-terminus of sE is fused to the trans-membrane and cytosolic tail of the type-I trans-membrane protein MHC-Iα for plasma membrane display of the protein, as previously reported (28). This assay has the potential to discriminate antibodies binding exclusively to dimeric E on the basis of the pH-dependent mobility of E protein: at pH7, that resembles the neutral extracellular environment, the protein is in a dimeric conformation but at pH6, mimicking the conformational changes that occur in the acidic endosome vesicles during infection, it moves to a pre-fusion monomeric conformation. When exposed to a neutral pH (pH7), E can physiologically dimerize, and therefore be recognized by the dimer specific monoclonal antibody EDE 1C10 (24). However, this interaction is completely abrogated when cells are exposed to a lower pH (pH6), due to the disruption of the dimer. Thus, with this dimer-specific antibody two populations of cells can be detected by flow cytometry – antibody-bound and unbound – depending on the assay conditions (Fig 3A - EDE). On the other hand, antibodies binding to epitopes that do not require dimeric conformation of the protein are not affected by the pH and therefore show no differences in the binding capacity, as shown using in-house made monoclonal antibody DIII-1B (Fig 3A – DIII-1B), that recognizes a linear epitope located on domain III (S1 Figure). This assay is primarily designed, and indeed works optimally, for monoclonal antibodies. Nevertheless, we reasoned that it would still be useful in evaluating the nature of antibodies in sera from vaccinees containing a mix of IgGs capable of binding to linear or conformational epitopes. As shown in Fig 3B, serum antibodies from sE-WT- and VLP-WT-immunized groups (first and third columns, respectively) seemed to not be particularly affected in the binding by the pH-dependent change of conformation. In contrast, sera from sE-cvD- and VLP-cvD-vaccinated animals showed a more consistent pH-dependent difference in the relative peak positions (Fig 3B; second and fourth columns, respectively).

**Figure 3:**
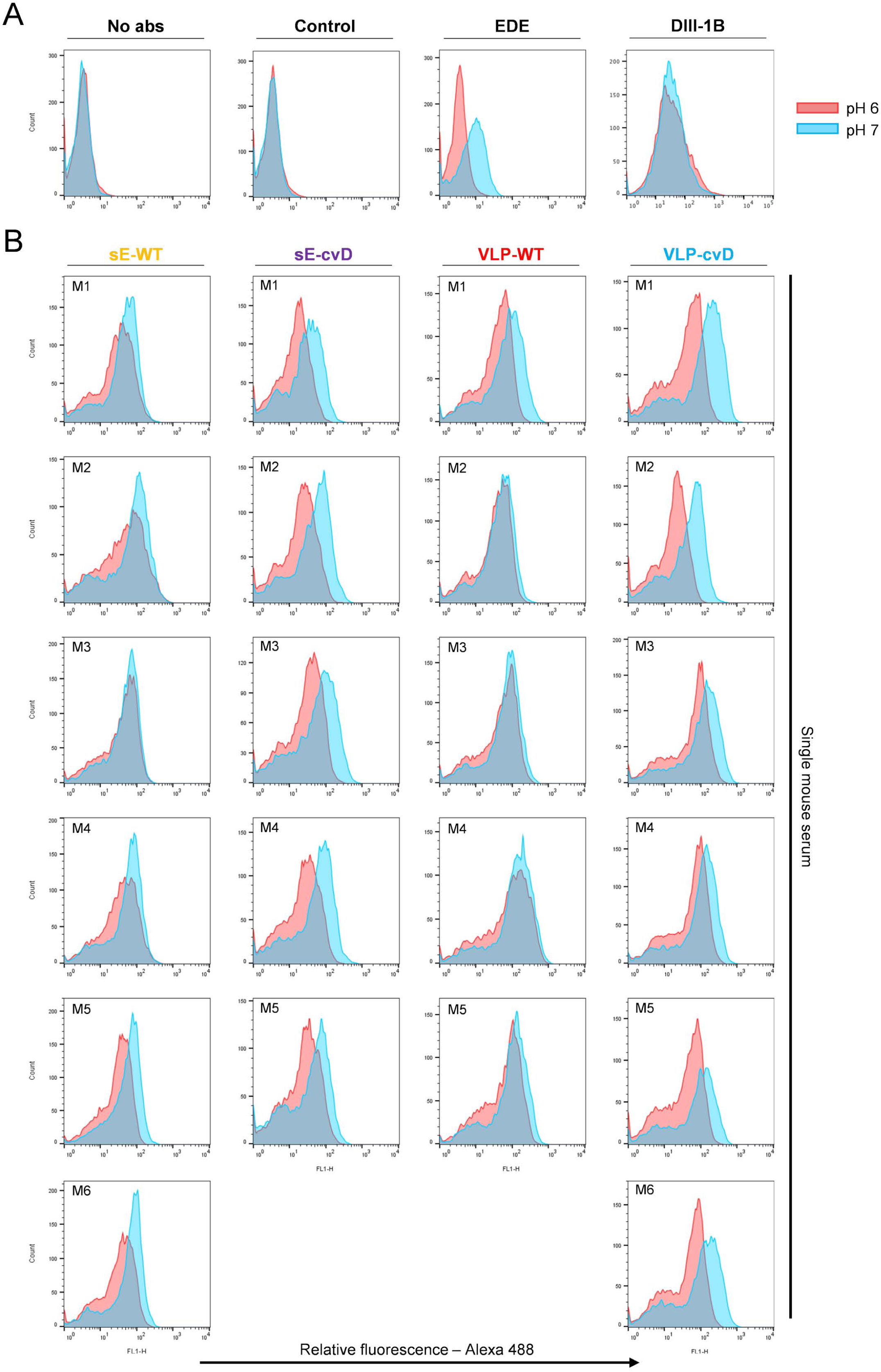
Determination of binding characteristics of serum IgGs to different sE conformations. Cells expressing sE on the cell surface were incubated with **(A)** secondary anti-mouse ALEXA 488 antibody only (No abs), pooled control sera (Control), monoclonal EDE antibody 1C10 (EDE), monoclonal DIII-1B antibody (DIII-1B) or **(B)** sera from immunised animals (M1 to M6) at pH6.0 (red) or pH7.0 (blue) as shown. Following washing, the bound antibodies were detected using a fluorescence-tagged secondary antibody and the relative fluorescence determined by flow cytometry using FACSCalibur. Data from three independent experiments were used

Taken together, the data suggest that cvD antigens elicited a population of antibodies more sensitive to changes in conformation then the one elicited by the WT immunisation.

### cvD antigens elicit neutralizing antibodies in mice

A second immunisation was performed using the same procedure shown in Fig 2A. Antibody titres were measured using exclusively the sE-cvD-BAP ELISA (Fig 4A). The neutralizing capacity of these sera was then determined in a micro-neutralization (MN) assay that we had previously developed (9). This sandwich ELISA accurately measures the levels of glycoprotein E in infected cells thus enabling quantitation of virus infectivity. Vero cells were infected with the Puerto Rican ZIKV strain PRVABC59 (an Asian lineage isolate) that had been pre-incubated for 1 hour with serially diluted mouse sera. Three days post-infection the level of cellular E protein was determined by the sandwich ELISA. Percentage of infectivity was calculated relative to E yield in cells infected in absence of sera. As shown in Fig 4B, antibodies elicited by VLP-WT or VLP-cvD neutralized virus infection significantly more strongly than their respective sE counterparts. Both the cvD antigens consistently produced higher (although not statistically significant) *in vitro* neutralization titres than their WT counterparts (Fig 4B). Sera from control group animals did not neutralize virus infectivity.

**Figure 4:**
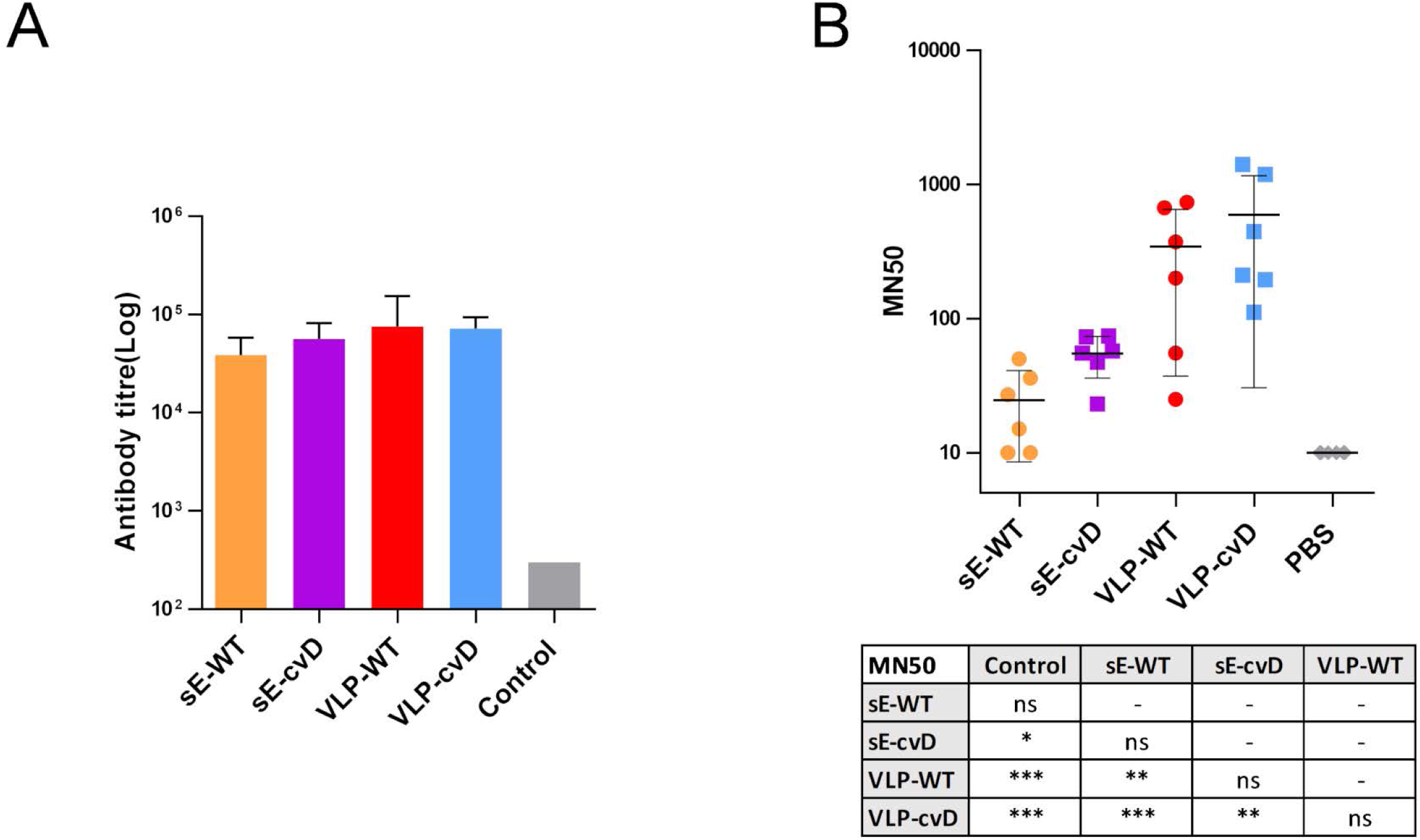
Determination of anti-E or neutralising antibody titres of sera from vaccinated animals. **(A) Anti-E titres:** ELISA plates coated with biotinylated dimeric E were incubated with serially diluted serum samples and the bound antibodies detected as described in Materials and Methods. Antibody titres were determined as described in the Figure 2 legend. (**B) Neutralisation of ZIKV infection:** Serially diluted samples of mouse sera were incubated with ZIKV for 1 hour before infecting Vero cells. At 72 hours post-infection, the intracellular levels of E were determined by capture sandwich ELISA and percentage of infectivity relative to the virus alone infection was calculated. The results were plotted as MN50 values - i.e. titres at which 50% neutralisation was achieved. Statistical analysis was performed using 2-sided ANOVA 95% confidence level with Tukey Pairwise comparison at 95% confidence (Minitab software). Data from three independent experiments were used

### cvD immunogens protect mice from ZIKV challenge

To assess *in vivo* efficacy of our candidate vaccines, we challenged a group of immunised animals with live virus. Analysis of their sera post-immunization confirmed the superiority of the cvD antigens in eliciting neutralizing antibodies (S2 Fig). Animals were challenged subcutaneously with 10^4^ pfu of ZIKV PRVABC59 after three immunisations with ZIKV antigens or PBS, as shown in Fig 5A. A scoring system was used to monitor the progress of the disease, based on severity of clinical signs and symptoms, as described in S1A Table. A score of 3 was considered as the humane endpoint. Animals were monitored for 9 days for their body weight changes (Fig 5B) and clinical signs (S1B Table). The PBS control group started losing weight at 4 days post-challenge (dpc) and subsequently exhibited clinical signs of infection and were euthanized at 7-8 dpc (Figs. 5B and C, grey lines). The sE-WT group lost less weight but exhibited clinical signs comparable to those seen in the PBS control group although at delayed onset (Figs. 5B and C, orange lines). One mouse of the sE-WT group succumbed to infection. A similar profile of weight change, clinical scores and survival were observed in VLP-WT group (Figs. 5B and C red lines) compared to the sE-WT group. Importantly, all animals immunised with cvD antigens survived the challenge, maintained a more stable weight profile and showed rapid recovery from the clinical signs of infection (Figs. 5B and C, purple and blue lines). Viremia was determined by RT-qPCR on blood samples taken at days 2, 3, 4 and 7 during the course of the challenge (Figs. 5D and E). Since the limit of the assay was determined as a titre of 10^2^ pfu equivalent/mL, for statistical analysis this value was given to all the samples that were below the limit of detection. As expected, PBS control mice showed very high viremia (>10^6^ pfu/mL) which peaked at 3 dpc. In contrast, in all vaccinated animals the viremia peaked at 4 dpc, although the levels varied. Specifically, the sE-WT-vaccinated animals displayed levels comparable to those observed in the PBS control group (>10^5^ pfu equivalent/mL). Instead, consistent reduction in viremia levels was observed in VLP-WT-, sE-cvD- and VLP-cvD-vaccinated animals which in the latter two groups was significant. In particular, the geometric mean of viral titre of 4×10^2^ pfu equivalent/mL was the lowest in VLP-cvD-vaccinated group. Relative organ viral load was analysed by RT-qPCR of viral RNA extracted from brain, spleen and sex organs, which were collected immediately after euthanasia (Fig 5F). ΔΔCT method was used to calculate the titre relative to an average of PBS control group. Although all four antigens reduced brain viral load, only cvD antigens reduced that of the sex organs. In case of spleen viral transmission, sE-cvD and VLP-cvD were better than sE-WT and control group whereas VLP-WT was only better than control group. All together, these data confirmed the unsuitability of wild-type antigens (especially sE-WT) whereas both cvD derivatives conferred full protection against ZIKV infection *in vivo*.

**Figure 5:**
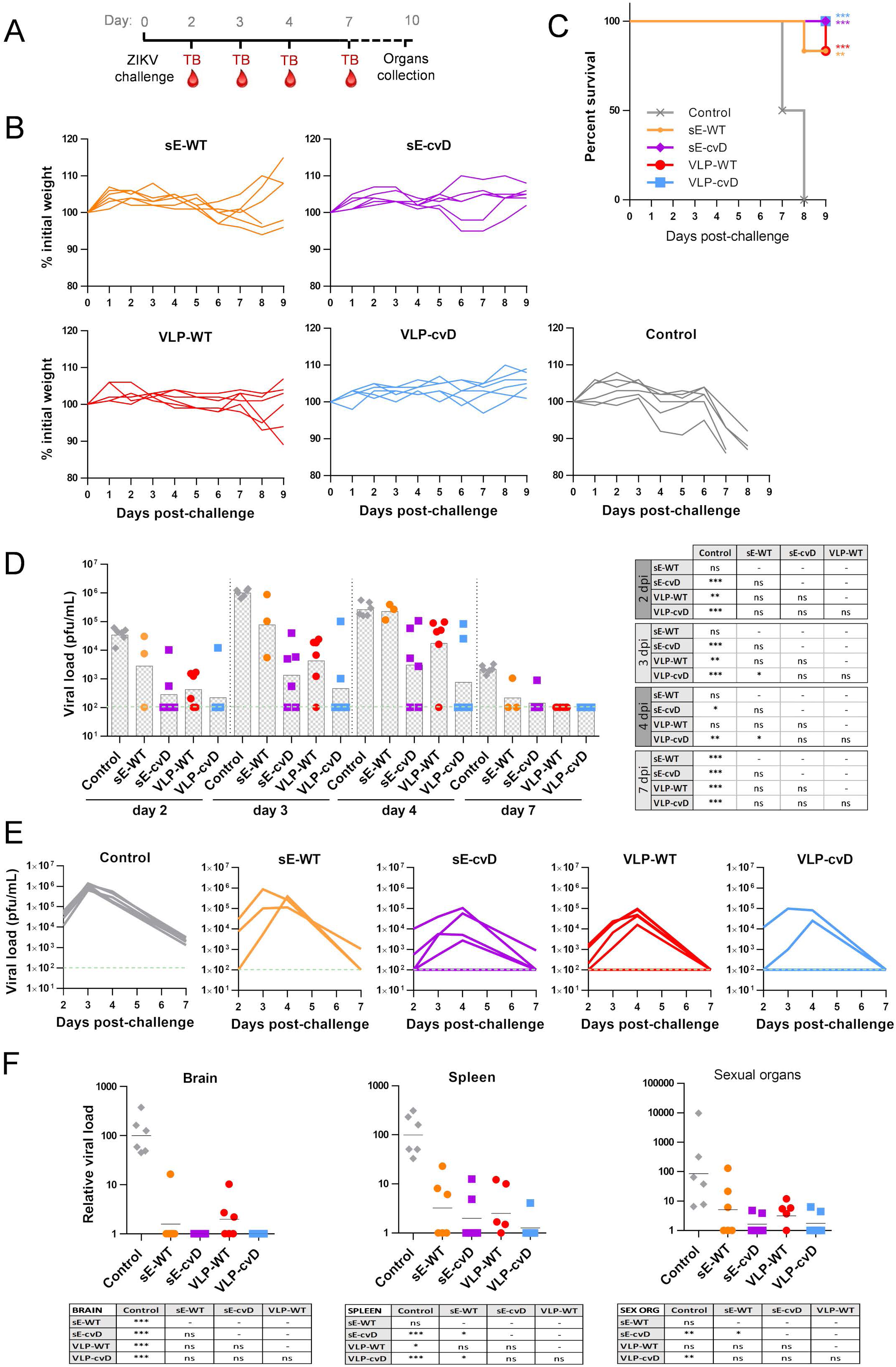
*In vivo* efficacy of candidate vaccines. (A) Schematic representation of the *in vivo* challenge protocol: Mice were challenged with 10^4^ pfu of ZIKV PRVABC59 one month after the primary immunisation and were monitored for up to 9 days. Test bleeds (TB) and organs were collected as shown. **Animals were weighed (B) and scored for clinical signs daily post-challenge with percentage of survival shown in (C).** Animals displaying a weight loss of 15% or more were euthanised. All the member of the control group reached the endpoint score 7-8 days post-challenge and were therefore euthanised. Statistical analysis was performed using Log-Rank (Mantel-Cox test) with GraphPad Prism software. (**D/E**) **The levels of ZIKV in the serum at day 2, 3, 4 and 7 post-infection were quantified by RT-qPCR and the results plotted as pfu/ml**. (D) The limit of detection was estimated to be 100 pfu/mL, indicated by the green line. Columns show mean of all mice. Statistical significance is reported in the table. **(F) Relative viral load in brain, spleen and sexual organs:** The presence of viral RNA in tissues was quantified by RT-qPCR and the results plotted as relative viral load calculated on the average of the PBS control group. Statistical significance is reported in the table. Statistical analysis was done using 2-sided ANOVA 95% confidence level with Tukey Pairwise comparison at 95% confidence with Minitab software.

### cvD reduces *in vitro* ADE

Due to the close relationship with DENV and other mosquito-borne flaviviruses, a ZIKV vaccine is very likely to elicit cross-reactive antibodies that may fail in neutralizing other flavivirus infection and instead lead to a worse disease outcome. Our candidate vaccines are designed in order to reduce this risk, limiting exposure of highly cross-reactive but low cross-neutralising epitopes in favour of broadly neutralizing antibodies. We performed an *in vitro* ADE assays using the K562 monocyte cell line that expresses the Fcγ-receptor. Infection of these cells occurs only in presence of antibodies opsonizing the virus and therefore mediating the entry through the Fcγ-receptor internalization. Viruses, pre-incubated with ten-fold serial dilutions of the sera were added to the cells, incubated for three days, and then analysed to determine the percentage of infection by cytofluorimetry. Experiment was performed in triplicate. (Fig 6).

**Figure 6:**
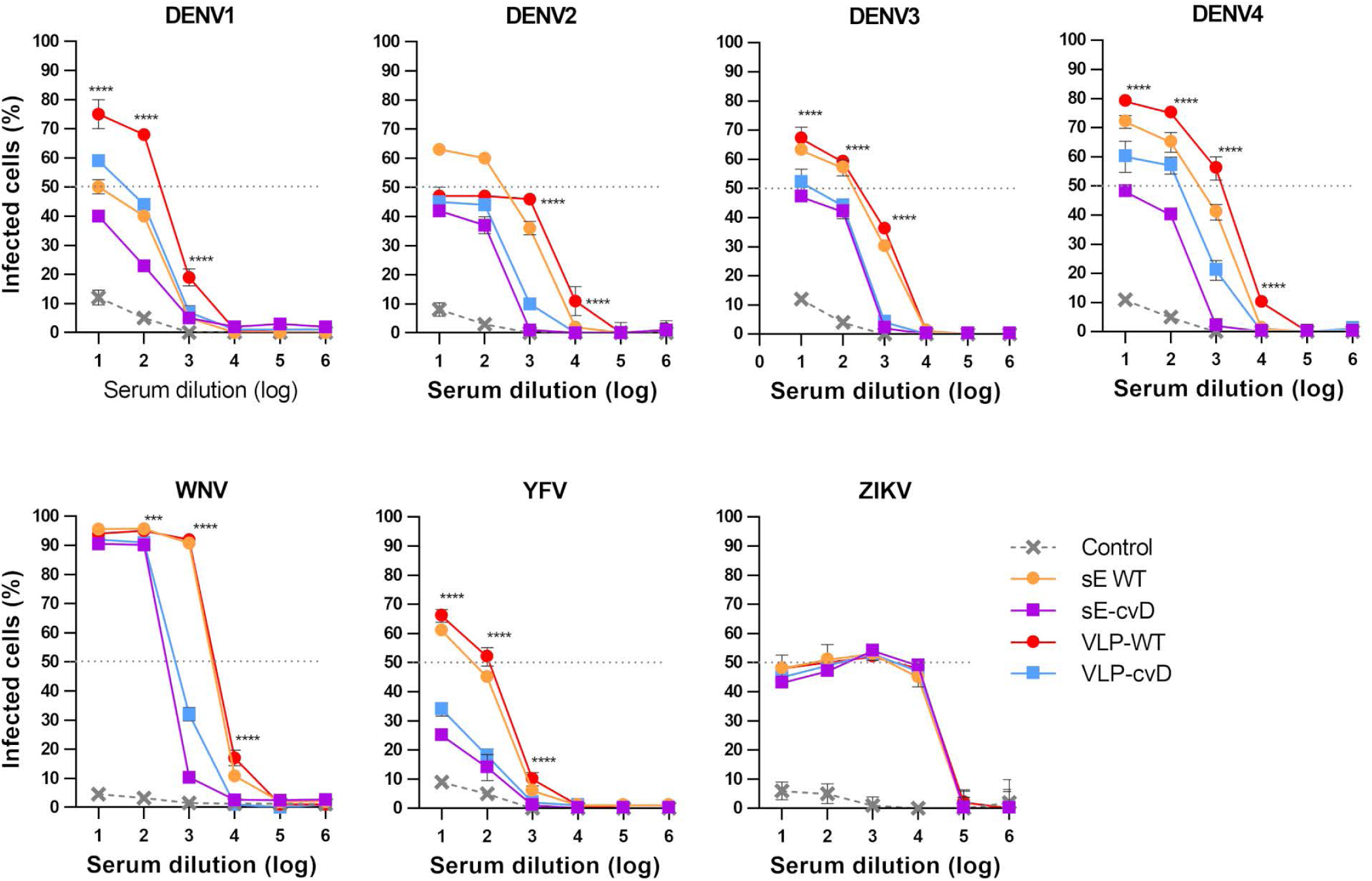
Effect of pooled sera on ADE of infection by all four DENV serotypes 1-4, ZIKV, WNV and YFV. Viruses were pre-incubated with 10-fold dilution of pooled sera for 1 hour before infecting K562 cells. Percentage of infected cells was calculated by cytofluorimetry. Statistical significance applies to the comparison between VLP-WT and sE-cvD/VLP-cvD. Statistical analysis was done using 2-sided ANOVA 95% confidence level with Tukey Pairwise multiple comparison (GraphPad software). Experiments were performed in triplicate.

We used ZIKV as a control in the assay and as expected all the sera gave the same pattern of infection, suggesting that the four groups of sera have a similar capacity to bind to ZIKV and therefore mediate infection. When tested against the four DENV serotypes, YFV and WNV, sera from the sE-WT group showed a percentage of infected cells between 50 and 100. Instead, 10 times lower levels of infection were observed with sE-cvD sera, suggesting a much lower level of cross-reacting antibodies. Similarly, sera from VLP-cvD-immunised mice exhibited a 10-fold reduced infectivity compared to sera from VLP-WT animals. Particularly interesting was the level of infected cells obtained after incubation with VLP-WT sera, which was significantly higher than the infection obtained with the sera from sE-cVD and VLP-cvD immunisations. These results suggest that the covalent dimer-based E vaccines (both, sE-cvD and VLP-cvD) confer a lower risk of ADE in comparison to their WT counterparts as determined by this experimental model.

### VLP-cvD protection coverage includes ZIKV African lineage

ZIKV has diverged decades ago into two lineages, the African and the Asian. Where the Asian lineage is responsible for the last epidemics and is linked to neurological outcomes and birth-defects, the African lineage is -intriguingly-well known to be more pathogenic in *in vivo* models (35). Since a safe ZIKV vaccine should guarantee coverage of both African and Asian lineages, we tested the VLP-cvD protectivity upon infection with a Ugandan (MP1751) isolate of ZIKV. Immunization and challenge were performed as previously described (Figs 2A and 5A). The PBS group lost weight starting from day 3 post-challenge and all mice reached the endpoint at day 6 (Figs 7A/B, grey lines. S2 Table). VLP-cvD-vaccinated group instead showed a stable body weight and all animal survived the challenge (Figs 7A and B, blue lines). PBS-immunised-control mice showed high peak of viremia (>10^7^ pfu/mL) at 4 dpc while the vaccinated mice showed a highly significant reduction in viremia (∼10^2^ pfu/mL) (Fig 7C). Also, virus dissemination to the brain was suppressed in the VLP-cvD immunized mice (Fig 7D). All together, these results confirm the broadly protective potential of our vaccine candidate.

**Figure 7:**
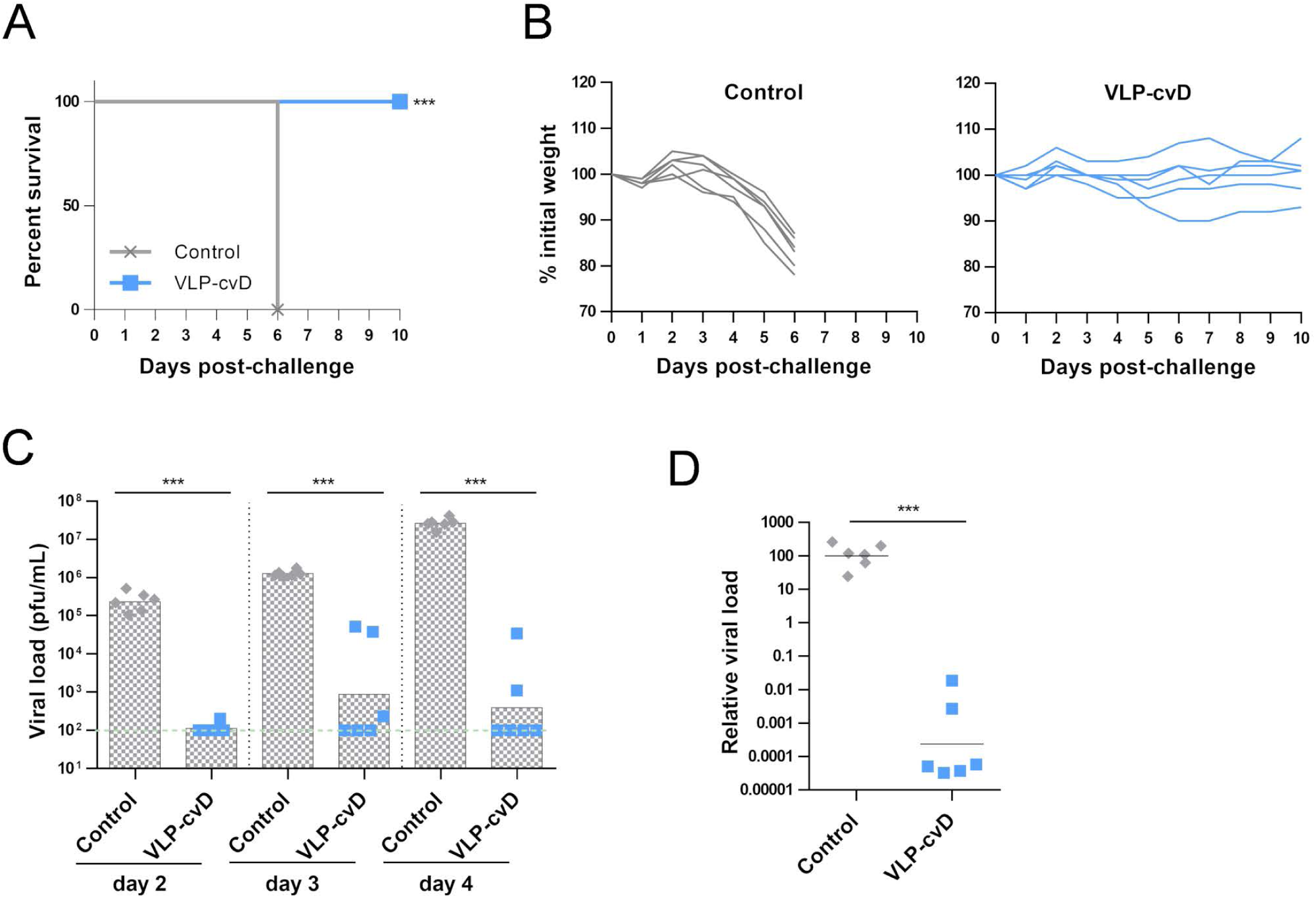
(A) Survival rate of vaccinated animals upon ZIKV challenge: Mice were challenged with 10^4^ pfu of ZIKV MP1751. All the member of the control group reached the endpoint score 6 days post-challenge and were therefore euthanised. Statistical analysis was performed using Log-Rank (Mantel-Cox test) with GraphPad Prism software. (**B) Body weight variations after challenge:** weight loss of mice after ZIKV infection. Animals showing a weight loss of 15% or higher were euthanised. (**C) Viral load in serum:** The presence of ZIKV in the serum at day 2, 3 and 4 post-infection was quantified by RT-qPCR. Green line indicates the limit of detection. **(F) Relative viral load in brain:** presence of viral RNA in tissues was quantified by RT-qPCR. Statistical analysis was done using 2 sided Two-Sample T-Test 95% confidence level with Minitab software.

## Discussion

Monoclonal antibodies recognizing quaternary epitopes that span on more than one E protein are reported to be the most neutralizing and cross-reactive class of antibodies (24, 36, 37). Nevertheless, they constitute only a minority of the antibodies elicited by a natural infection, where the larger response focuses on poorly neutralizing and cross-reactive epitopes located on DI/DII (15, 38). The incomplete maturation of particles and the mobility of E reduce their exposure, especially to dimeric epitopes, in favour of the Fusion-Loop epitope. However, our vaccines are designed in order to lock E in a dimeric conformation, impairing its disassembly and forcing the protein to display the desired epitopes.

E-homodimer stability was proved several times to be affected by temperature: dimerization is favoured at 28°C and reduced at 37°C, hence physiological temperature is another element that contributes to impair dimers exposure to the immune system (39). But 28°C is also a crucial temperature for expression and secretion of E protein, both in the soluble form but also as part of VLPs (28). Since this temperature is not compatible with genetic vaccination approaches, in which DNA or RNA encoding the antigen is administered and the antigen is then produced by the host cells at physiological temperature, the full potential of our antigens as candidate vaccines was evaluated in a protein-based vaccination approach.

Recently, a characterization of two dimeric E antigens was published, dimerization was achieved by the A264C mutation or replacing the E transmembrane domain with the FC fragment of a human IgG (40). While this manuscript was under preparation, two other articles reported development and evaluation of dimer-based subunit vaccines similar to that described in here (41, 42). The authors showed that in all three cases the antigens were able to induce in mice the production of neutralizing antibodies. However, our work is the first application of E covalent dimers to virus-like particles that proved in our hands to be a vaccine candidate superior to the soluble dimer.

We tested the potential of E covalent-dimers when expressed in the form of soluble protein, lacking the stem-anchor, and also as VLP. In comparison with sE-WT we observed a dramatic effect of the cvD mutation on the immune response, with a drastic reduction of antibodies binding to monomeric E and an impressive increase in protective activity upon lethal challenge. When the comparison was performed with the VLPs, the effect was less dramatic in terms of anti-dimer antibody titres. This is likely due to the thermal stability of dimers that is higher in ZIKV than in DENV (43), ZIKV E dimers are more stable, as already proven by the necessity of a double disulphide bridge to lock DENV E dimer when one bridge is sufficient in ZIKV E. In addition, our ELISA and binding assays are based on E subunits presented in a monomeric or dimeric form but cannot fully recreate the complex symmetry of E protein on the viral particles therefore cannot quantify the contribution of antibodies binding to adjacent dimers. However, the difference in antibody response and protectivity was enough to achieve an important reduction in DENV ADE. In this regard it is noteworthy that the VLP-cvD lacked prM indicating their complete maturation. In contrast, VLP-WT contained prM that can elicit anti-prM antibodies upon vaccination. The prM protein is naturally present in DENV or ZIKV particles, due to incomplete maturation, but prM antibodies from DENV patients showed no or poor neutralizing activity and may instead likely induce ADE (44). All together these observations strongly suggest that covalently linked E dimer can bring even higher benefits to the development of a DENV vaccine.

The development of a ZIKV vaccine requires attention to the possible cross-reaction with DENV. ADE of infection between different DENV serotypes is widely recognised as the cause of dengue shock syndrome (DSS) but the role that infection may play at any point in influencing DENV infections is still unclear. So far animal models have not been able to replicate the full repertoire of the antibody response in humans. In addition, it is difficult to reproduce severe DENV infection in animal models; in most cases the severity of infection is based on increased viremia in the infected animals. *In vitro* tests performed with DENV-positive sera or DENV monoclonal antibodies showed cross-reactivity and ADE of ZIKV infection, similar to experiments performed in mouse models (20, 45-47). In other cases analysis performed in non-human primate models ruled out a negative effect of previous DENV exposure on ZIKV infection (48), which was later confirmed by population studies with asymptomatic ZIKV infection in subjects positive for DENV antibodies (49). Concerns of cross-reaction and ADE were also raised about vaccination against other flaviviruses. *In vitro* studies supported negligible risk of ADE of ZIKV after tick-borne encephalitis virus vaccination and no clinical evidence of increased disease severity in vaccinated people has emerged so far (50). The fear of predisposing vaccinated individual to DSS generated a reluctance to deploy the YFV vaccine in DENV endemic areas, but a recent long-term study showed no evidence of increased risk (51). However, the sequence homology between ZIKV and DENV is high with elevated antibody cross-reactivity (15, 20) and how ZIKV vaccination can affect DENV pathogenesis is a pertinent question.

We evaluated the ADE potential (on DENV, YFV and WNV) of the different sera using K562 cells, which by expressing high levels of FcγRIIA, significantly favours FcγR-mediated infection, making these cells particularly prone for ADE-mediated infection. In contrast to sera of sE-WT vaccinated animals that showed high level of cross-reaction, ADE was strongly reduced when E was locked in the dimeric form (sE-cvD) and properly folded, likely because of the exposure of conserved and poorly neutralising epitopes on DI/DII. The most interesting results were provided by the comparison between WT and cvD VLPs. Such containing un-mutated E, despite being the major candidate exploited so far and being able to sufficiently protect mice from lethal infection, induced *in vitro* ADE at a level comparable to sE-WT, or even higher as in the case of DENV1 and DENV4. This raises safety concerns about a vaccine that, despite protecting from ZIKV infection, may bring more adverse effects on subsequent DENV infections. Once again, this risk is reduced with the VLP-cvD antigen. However, to what extent ADE *in vitro* mimics any *in vivo* effects remains to be determined.

ADE of DENV upon ZIKV immunisation is a risk that cannot be underestimated, and the development of engineered, safe vaccine is probably an essential requirement to tackle this concerning public health challenge. Our two immunogens proved the high potential of engineered E protein locked in a dimeric conformation as a suitable vaccine candidate, with the most promising results achieved when the protein is part of a structurally more complex antigen presented in the form of a virus-like particle.

## Material and Methods

### Cell lines and virus strains

Expi293F (Thermo Fisher Scientific) embryonic human kidney adapted to serum-free conditions) cells were grown in Expi293™ Expression Medium as per the manufacturers’ protocol. Vero E6 cells were grown in Dulbecco’s Modified Eagle’s Medium (DMEM) (Life Technologies) containing 10% fetal bovine serum (FBS) (Life Technologies) and penicillin-streptomycin (Gibco), K562 cells were grown in Roswell Park Memorial Institute (RPMI) 1640 medium (Life Technologies) containing 10% Fetal Bovine Serum (FBS) (Life Technologies). ZIKV PRVABC59 (kindly supplied by BEI Resources; Accession Number KX087101) and ZIKV MP1751 (005V-02871; kindly supplied by Public Health England: Accession Number KY288905.1) were used for infection experiments, micro-neutralisation and animal challenge.

### Plasmid DNA constructs

ZIKV sE-encoding sequence (codon 1-404) was amplified from ArD158095 strain (Accession number KF383121.1) as described in Sloan Campos et al^(28)^. sE fused to a N-terminal immunoglobulin leader sequence (sec) and a C-terminal V5 tag (GKPIPNPLLFLD) was cloned into a pVax vector. A mammalian codon-optimized ZIKV prME gene sequence, flanked by the C-terminal portion of C and the N-terminal reside of NS1, was obtained from ZIKV PE243 Brazilian strain (Accession number KX197192.1^(52)^) (aa 105-815 of the polyprotein) and cloned into a pDIs vector. The A264C mutation was introduced by site-directed mutagenesis into both plasmids.

### Protein expression and purification

sE-WT, sE-cvD, VLP-WT and VLP-cvD were expressed using ExpiFectamine™ 293 Transfection Kit (Thermo Fisher Scientific) following manufacturer’s instructions. After 16 hours, cells were moved to 28°C. At 5 days post-transfection the supernatant was harvested and filtered. sE proteins were purified using the V5-tagged Protein Purification Gel (Caltag Medsystems Ltd) eluting with 2 mg/mL of V5 peptide. VLPs, they were pelleted down by ultracentrifugation (115,000 g, 4°C, 2 hours) (Sorvall discovery 90SE with Surespin630 rotor) through a cushion of 20% sucrose in TN Buffer (20 mM Tris and 120 mM NaCl). The pellet was re-suspended in TN buffer and loaded on discontinuous density gradient made by sodium potassium tartrate and glycerol in TN buffer (29). Tartrate concentration ranged from 10 to 30% with interval of 5% whilst that of glycerol ranged from 7.5 to 22.5% with interval of 3.75%. After centrifugation (Sorvall discovery 90SE with TH641 rotor) at 175,000 g, 4°C, 2 hours, fractions were collected and analysed for the presence of ZIKV E by western lot. ZIKV E protein-positive fractions were pooled, dialysed against Dulbecco’s phosphate-buffered saline (DPBS) (Life Technologies) and concentrated using spin column (Amicon® Ultra-15 (100 kDa), Merck Millipore) before being subjected to size-exclusion chromatography. Briefly, ∼500 µL of concentrated pooled fractions was loaded onto HiPrep 16/60 Sephacryl S-500 HR column (GE Healthcare) then 1.5 column volume of mobile phase (DPBS) was run through the column at flow rate of 0.5 mL/min using the AKTA Pure (GE Healthcare) system. Fractions were collected and tested for ZIKV E protein. Positive fractions were pooled and concentrated using the Amicon® Ultra-15 (100 kDa; Merck Millipore) spin column. The concentration of the purified proteins was determined using NanodropOne (ThermoScietific).

### SDS-PAGE and western blot

sE samples were subjected to 10% SDS-PAGE, and the fractionated proteins detected by direct staining of the gel with InstantBlue (Sigma) or by western blot. VLP samples were separated by 10% or 14% SDS-PAGE later blotted to PVDF membrane (Immobilon®-FL, Merck Millipore) and blocked overnight with ODYSSEY® blocking buffer, LI-COR then incubated DIII1B antibody (anti-ZIKV E DIII generated in-house as described in S1 Figure) or ZIKA prM antibody (GeneTex) for 1 hour followed by anti-mouse IgG (IRDye® 800CW, LI-COR) and anti-rabbit IgG (IRDye® 680RD, LI-COR). Images were acquired by LICOR machine.

### Electron microscopy

VLPs were adsorbed for 3 min to Formvar carbon films mounted on 400 mesh per inch copper grids (Agar Scientific). Samples were washed three times with distilled water and stained with 2% saturated uranylacetate (Agar Scientific) for 2 min at room temperature. Specimens were analysed in a transmission electron microscope (JEM-1200 EX II, JEOL) equipped with a CCD camera (Orius, Gatan) at an acceleration voltage of 80 kV.

### Animal immunisation

Four-week old male and female *Ifnar1*^*-/-*^ mice (A129, 129S7 background; Marshall BioResources) (n=6) were subcutaneously immunised with ZIKV antigen formulated in aluminium hydroxide gel (1% ALUM, Brenntag) combined with 5 µg monophosphoryl lipid A (MPLA)(InvivoGen) or PBS containing the adjuvant. Purified sE antigens used in each immunisation contained 10 µg protein whilst it was 2 µg in case of VLPs. Mice were immunised at 0, 2 and 3 weeks and bled 4 weeks after primary immunisation for antibody titration and micro-neutralisation assay. Four weeks after primary immunisation, mice were challenged subcutaneously with 10^4^ pfu of Puerto Rican ZIKV (PRVABC59) or Uganda ZIKV (MP1751). Blood was collected at 2, 3, 4 and 7 dpc and 10 µL of sera were assessed by RT-qPCR. Mice were euthanised when they exhibited three or more signs of moderate severity or lost more than 15% body weight, otherwise 9/10 days after challenge.

### Animal Ethics

All animal research described in this study was approved by the University of Glasgow Animal Welfare and Ethical Board and was carried out under United Kingdom Home Office Licenses, P9722FD8E, in accordance with the approved guidelines and under the UK Home Office Animals (Scientific Procedures) Act 1986 (ASPA).

### ELISA

Recombinant biotinylated proteins (sE, sE-cvD and DIII) were expressed at 28°C using ExpiFectamine™ 293 Transfection Kit (Thermo Fisher Scientific). Cell supernatant was harvested and dialyzed. Biotinylated proteins were captured in ELISA plates pre-coated with 5 µg/mL of Avidin (Sigma) in Na2CO3/NaHCO3 buffer pH 9.6, subsequently blocked with PBS containing 0.05% Tween-20 and 1% bovine serum albumin (BSA-Sigma). Serial dilutions of mouse sera were tested for binding to the biotinylated proteins and the bound antibodies detected using HRP-conjugated anti-mouse IgG A4416 (Sigma) and TMB substrate (Life Technologies).

### Antibody binding assay

This assay was performed as previously described in (paper in submission). HEK cells stably expressing ZIKV sE protein on the surface were blocked in 1% BSA in PBS at pH 6 or 7 and then incubated with mouse sera diluted 1:500 in the same solution. After wash, cells were incubated with secondary anti-mouse Alexa 488 (Jackson Immunoresearch) 1:50000 in 1% BSA PBS pH 7 and analysed by cytofluorimetry in a FACSCalibur (BD Biosciences).

### Micro-neutralisation (MN) assay

This assay was performed as described in Lopez-Camacho et al (9). Briefly, 7 × 10^3^/well of Vero cells were seeded in 96-well plates and incubated at 37 °C in 5% CO2. Next day, three-fold serially diluted mice sera were first incubated at 37°C for 1 hour with 100 pfu/well ZIKV strain PRVABC57. The serum/virus mix was then used to infect cells. After 1 hour of incubation at 37 °C, 100 µL of medium was added to each well. At day 3 post-infection, cells were lysed in lysis buffer (20 mM Tris-HCl [pH 7.4], 20 mM iodoacetamide, 150 mM NaCl, 1 mM EDTA, 0.5% Triton X-100 and Complete protease inhibitors) and the viral E protein quantitated by sandwich ELISA (see below). The amount of E protein detected correlates with the level of virus infectivity which was presented as % of ZIKV infectivity relative to the control (i.e., virus not pre-incubated with immune sera). The MN50 titre was defined as the serum dilution that neutralised ZIKV infection by 50%.

### Sandwich ELISA to assess ZIKV infectivity

ELISA plates were coated with 3 μg/mL of purified pan-flavivirus MAb D1-4G2-4-15 (ATCC® HB112TM) in PBS and incubated overnight at RT and subsequently blocked for 2 hours at RT with PBS containing 0.05% Tween-20 and 2% skimmed milk powder. After washing with PBST, ZIKV-infected cell lysates were added and incubated for 1 hour at RT. Wells were washed with PBST, incubated with anti-ZIKV E polyclonal R34 IgG (9) at 6 μg/ml in PBST for 1 hour at RT and washed again. Antibodies bound to ZIKV envelope protein were detected using HRP conjugate anti-rabbit IgG 7090 (Abcam) and TMB substrate (Life Technologies).

### Quantitation of viral RNA by RT-qPCR

Viral RNA was extracted from 10 µL of mouse sera using QIAamp® viral RNA Mini Kit (Qiagen) or about 20 mg of organs homogenised by Precellys Lysing Kit Hard tissue grinding (Bertin Technologies) using RNeasy viral mini kit. The viral load was measured by RT-qPCR using One-Step SYBR® Primescript™ RT-PCR kit II (Takara). CT values of serum samples were used to calculate serum viral titre according to regression equation built by RNA extracted from 10 µL of 10^2^-10^6^ pfu/mL of ZIKV (PRVABC59 or MP1751). In case of relative organ viral load, CT values of ZIKV gene and internal control B2M gene were used for calculating ΔCT values. Average ΔCT of PBS-injected mice was used as reference to calculate ΔΔCT value. Primers pair for PRVABC59 ZIKV gene was Forward:5’-GTTGTCGCTGCTGAAATGGA-3’ and Reverse:5’-GGGGACTCTGATTGGCTGTA-3’. Primers pair for MP1751 ZIKV gene was Forward:5’-ACTTCCGGTGCGTTACATGA-3’ and Reverse:5’-GGGCTTCATCCATGATGTAG-3’. Primers pair for the B2M genes was Forward: 5’-CGGCCTGTATGCTATCCAGA-3’ Reverse: 5’-GGGTGAATTCAGTGTGAGCC -3’.

### ADE assay

Ten-fold serial dilutions of pooled sera were mixed with 4×10^3^ pfu of each virus and incubated for 1.5 hour at 37 °C before mixing with 4×10^4^ K562 cells. After incubation at 37°C for 2 days (for WNV) or 3 days (all other viruses), cells were fixed with 2% PFA for 30 min and then washed in PBS. Blocking and permeabilization buffer (0.1% saponine, 2 % FBS, 0.1% NaN3 in PBS) was added to cell for 30 min at 4 °C. Cells are incubated with mAb 4G2 (1 µg/mL) for 1hour at 4 °C followed by secondary anti-mouse Alexa 488 (Jackson Immunoresearch, 1:50000). After washing with PBS, cells are re-suspended in blocking buffer without saponine and analysed by cytofluorimetry in a FACSCalibur (BD Biosciences).

### Statistical analysis

Normality was determined by Ryan-Joiner Normality test with Minitab Software. Statistical analysis was done as indicated in figure legends, with Minitab or GraphPad Prims softwares. *p<0.05, **p<0.01, ***p<0.001.

## Acknowledgment

We thank Marion McElwee and James Streetley for help with imaging VLPs by electron microscopy. We acknowledge the provision of ZIKV strains PRVABC59 and MP1751 (005V-02871) by BEI Resources and Public Health England/EVAg, respectively. This research is funded by the Department of Health and Social Care using UK Aid funding and is managed by the NIHR (AHP and AK); the UK Medical Research Council (MC_UU12014/2 and MC_UU_12014/8) (AHP and AK). The views expressed in this publication are those of the author(s) and not necessarily those of the Department of Health and Social Care. This project was also partially funded through the European Union’s Horizon 2020 research and innovation programme under ZikaPLAN grant agreement No 734584 (JME). The funders had no role in study design, data collection and analysis, decision to publish, or preparation of the manuscript.

## Author contribution

The CVR has adopted the CRediT taxonomy (https://casrai.org/credit/). Authors’ contribution is as follows. GDL: conceptualization, data curation, formal analysis, investigation, project administration, methodology, supervision, validation, visualization, writing - original draft, writing - review & editing; RT: data curation, formal analysis, investigation, validation, visualization, writing – original draft, writing review & editing; JD: data curation, investigation, writing - review & editing; CS: investigation, visualization, writing - review & editing; MP: investigation, methodology, validation, writing - review & editing; RS: investigation, writing - review & editing; HES: resources, writing - review & editing; JME: resources, writing - review & editing; AK: funding acquisition, resources, writing - review & editing; JB: resources, writing - review & editing; ORB: conceptualization, resources, writing - original draft, writing - review & editing, AHP: conceptualization, funding acquisition, project administration, resources, supervision, writing - original draft, writing - review & editing.

## Legends to Supplementary Figures and Tables

**S1 Figure:**
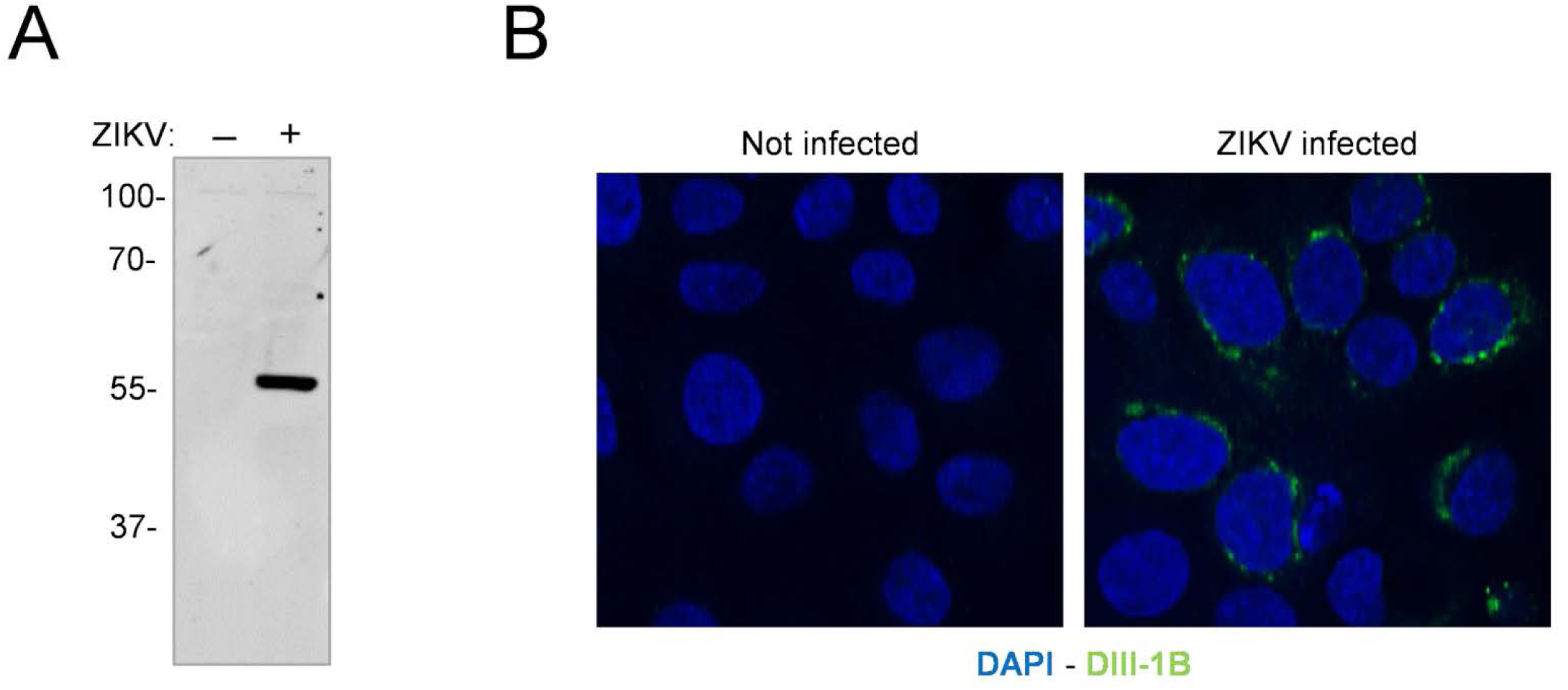
Characterisation of the mouse monoclonal antibody (MAb) DIII-1B. MAb DIII-1B was obtained using standard hybridomas technology from Balb/c mice that were immunised with recombinant domain III of ZIKV E protein. The specificity of MAb DIII-1B was tested by **(A)** western immunoblotting of VERO cells that were mock-infected (-) or infected with ZIKV (+). As expected, MAb DIII-1B specifically bound to ZIKV E protein. Protein molecular weight ladder is shown on the left in kDa. Separately, the binding specificity of MAb DIII-1B was also tested by **(B)** indirect immunofluorescence of uninfected or ZIKV-infected A549-NPro cells. Green signal indicates antibody binding to ZIKV E protein. Cell nuclei were stained with DAPI (blue).

**S2 Figure:**
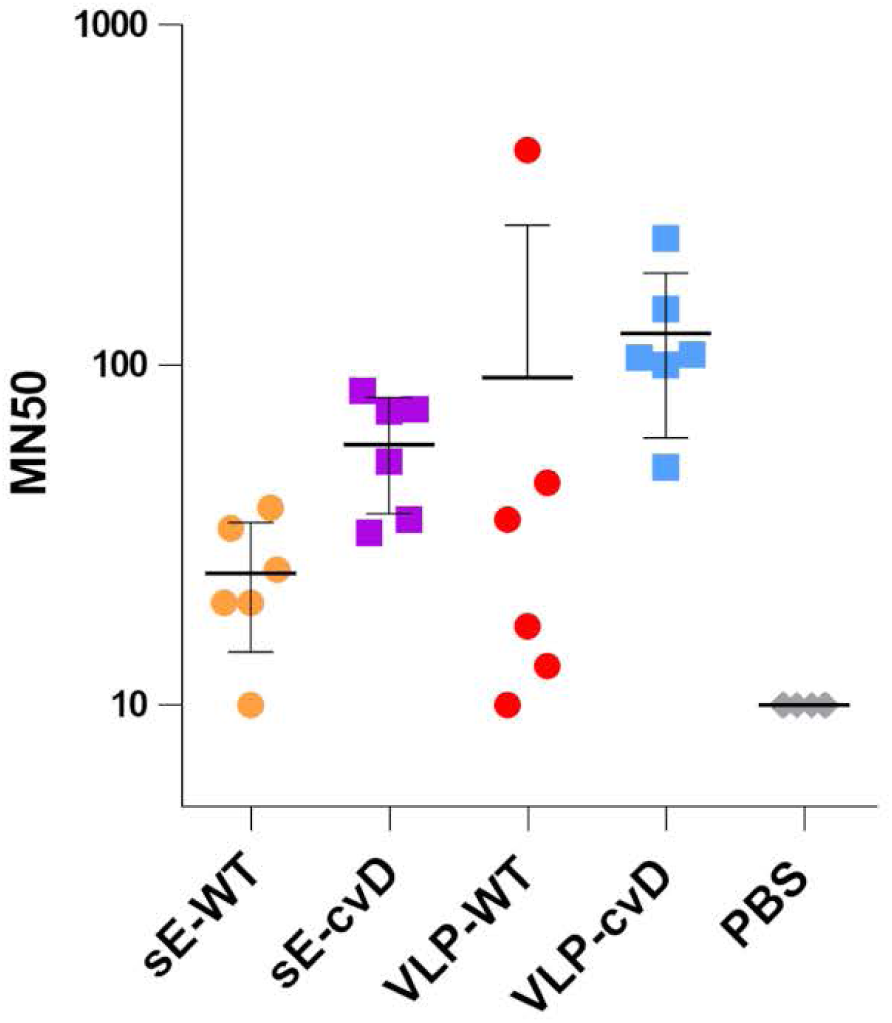
Determination of neutralising antibody titres of sera from vaccinated animals. Serially diluted samples of mouse sera were incubated with ZIKV for 1 hour before infecting Vero cells. At 72 hours post-infection, the intracellular levels of E were determined by capture sandwich ELISA and percentage of infectivity relative to the virus alone infection was calculated. The results were plotted as MN50 values - i.e. titres at which 50% neutralisation was achieved.

**Supplementary Table 1:**
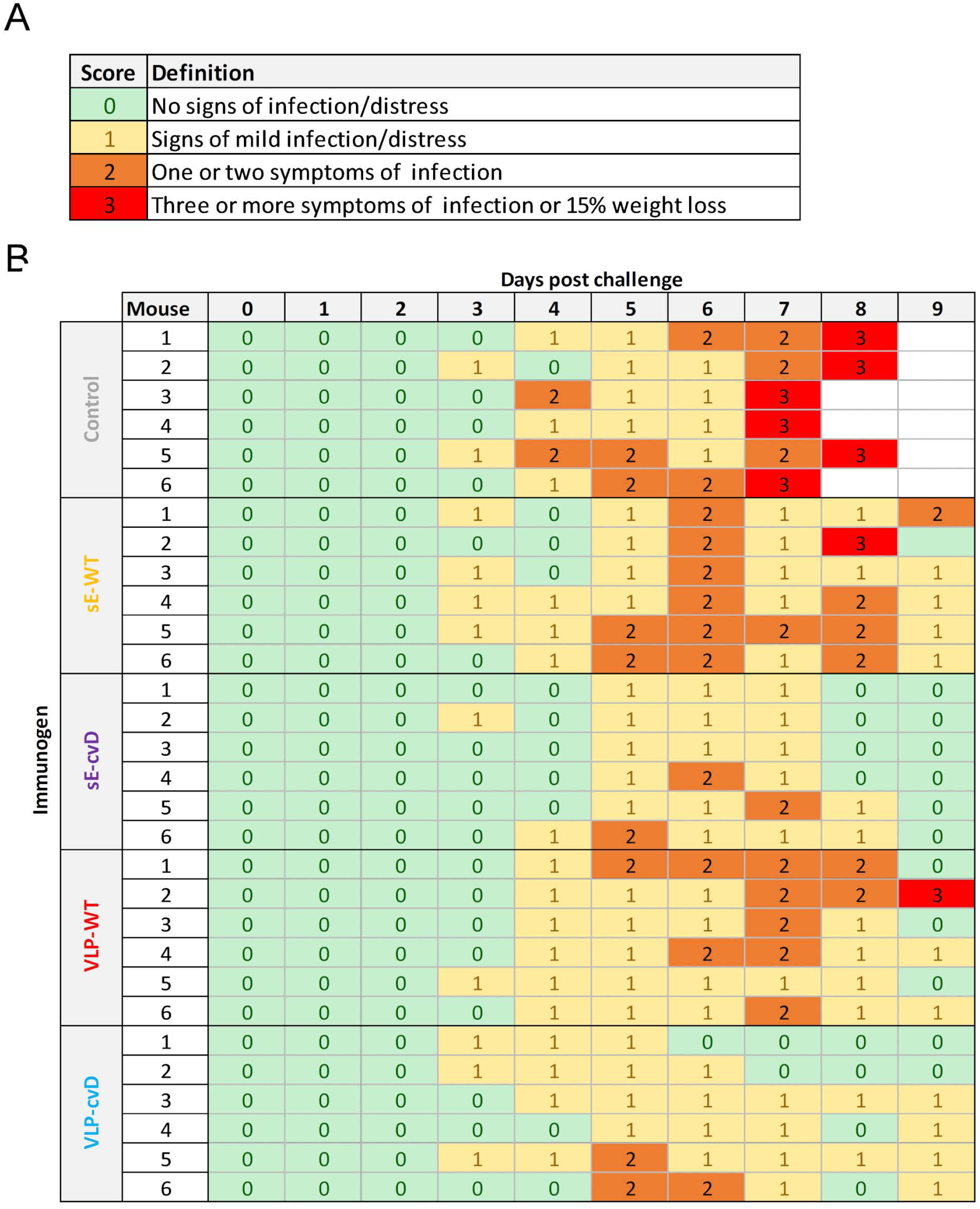
**(A)** Legend of scoring system used to monitor animal health following ZIKV challenge. **(B)** Table showing the score attributed to each animal after ZIKV PRVABC59 challenge.

**Supplementary Table 2:**
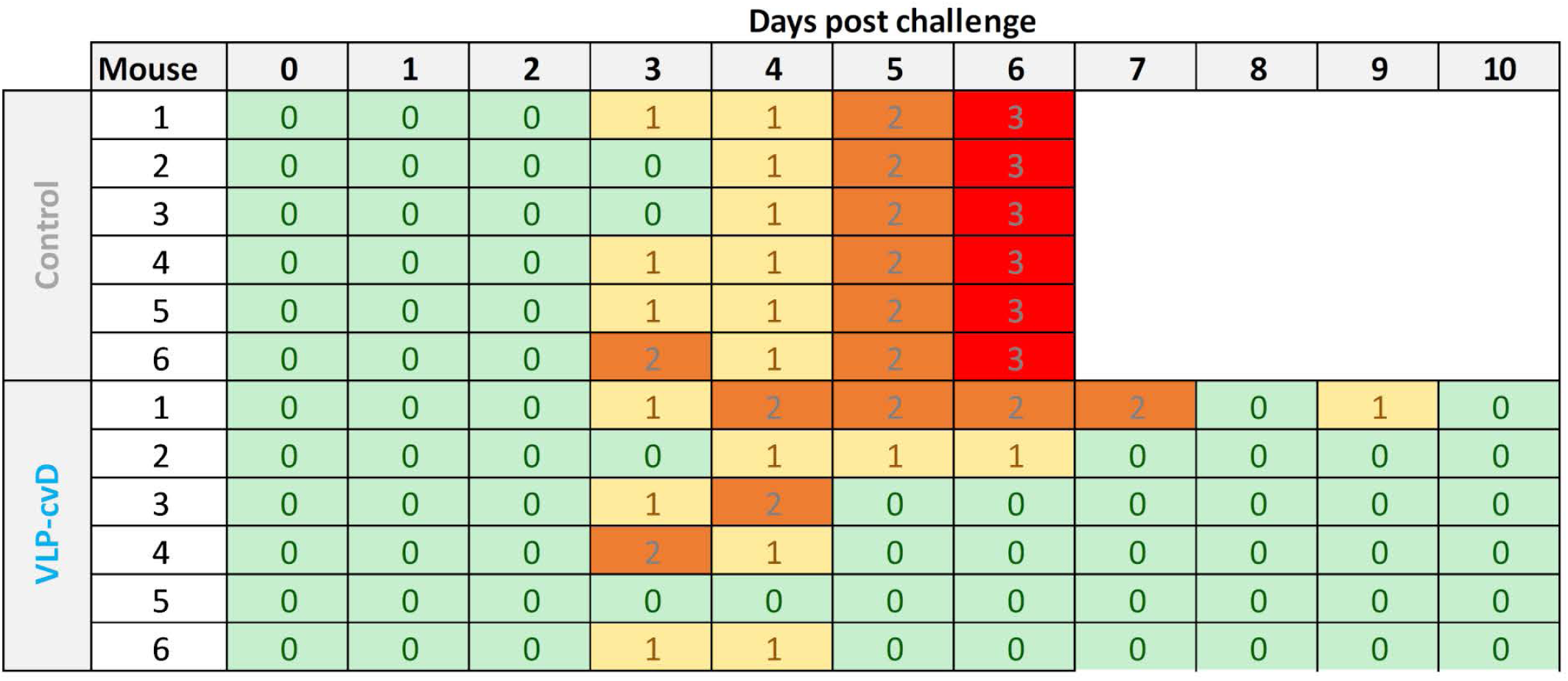
Clinical scores attributed to each animal after ZIKV MP1751 challenge.

